# Detection of *Neisseria meningitidis* in Saliva and Oropharyngeal Samples from College Students

**DOI:** 10.1101/2021.06.09.447670

**Authors:** Willem R. Miellet, Rob Mariman, Gerlinde Pluister, Lieke J. de Jong, Ivo Grift, Stijn Wijkstra, Elske van Logchem, Janieke van Veldhuizen, Marie-Monique Immink, Alienke J. Wijmenga-Monsuur, Nynke Y. Rots, Elisabeth A.M. Sanders, Thijs Bosch, Krzysztof Trzciński

## Abstract

**Objectives:** Since conjugated polysaccharide vaccines reduce carriage of vaccine-type *Neisseria meningitidis* strains, meningococcal carriage is an accepted endpoint in monitoring vaccine effects. We have assessed vaccine-type genogroup carriage prevalence in students at the time of MenACWY vaccine introduction in The Netherlands. In addition, we evaluated the feasibility of saliva sampling and qPCR-based detection method for the surveillance of meningococcal carriage.

**Methods:** Paired saliva and oropharyngeal samples, collected from 299 students, were cultured for meningococcus. The DNA extracted from all bacterial growth was subjected to qPCRs quantifying meningococcal presence and genogroup-specific genes. Samples negative by culture yet positive for qPCR were cultured again for meningococcus. Results for saliva were compared with oropharyngeal samples.

**Results:** Altogether 74 (25% of 299) students were identified as meningococcal carrier by any method used. Sixty-one students (20%) were identified as carriers with qPCR. The difference between number of qPCR-positive oropharyngeal (n=59) and saliva (n=52) samples was not significant (McNemar’s test, *p*=0.07). Meningococci were cultured from 72 students (24%), with a significantly higher (*p*<0.001) number of oropharyngeal (n=70) compared with saliva (n=54) samples. The prevalence of genogroups A, B, C, W, and Y was none, 9%, 1% and 6%, respectively, and 8% of students carried MenACWY vaccine-type genogroup meningococci.

**Conclusions:** We show that the detected prevalence of meningococcal carriage between oropharyngeal and saliva samples was nondifferent with qPCR and moreover, detection with both samples was highly concordant. Saliva is easy to collect and when combined with qPCR detection can be considered for meningococcal carriage studies.

## INTRODUCTION

*Neisseria meningitidis* (meningococcus) is a commensal of the human upper respiratory tract (URT) and a major cause of invasive bacterial disease [1]. Adolescents are at increased risk of invasive meningococcal disease (IMD) [2]. Following an outbreak of serogroup W IMD in the Netherlands in the fall of 2018, a monovalent conjugate polysaccharide vaccine targeting serogroup C (NeisVac-C, Pfizer) was replaced in the National Immunization Program with a tetravalent conjugated polysaccharide vaccine (Nimenrix, GlaxoSmithKline) targeting serogroups C, A, W, and Y [3]. Initially, the MenACWY vaccine was given only to 14-months-old children but since 2019 it is also offered to 14 year olds [4]. Conjugated vaccines not only protect against disease but also reduce carriage of vaccine-type (VT) strains [5]. Since the prevalence of meningococcal carriage is reported to peak in adolescents and young adults, vaccination in teenagehood is expected to induce herd protection across the population [2]. Effects of conjugated polysaccharide vaccines can be monitored via surveillance of carriage [6]. For this, reliable and efficient detection methods for meningococcus are required.

Oropharyngeal samples have been widely used to detect meningococcal carriage as it has been reported that oropharyngeal samples are more sensitive than nasal or nasopharyngeal samples [7]. While a role for saliva in meningococcal transmission has been implicated in multiple studies [8–16], and closely-related Neisseria species are often cultured from saliva [17], few studies have tested saliva for meningococci [18–21]. In general, saliva is described to be poorly suited for meningococcal detection [19]. Unlike oropharyngeal and nasopharyngeal swabs, saliva sampling is noninvasive, and oral fluids can be easily self-collected.

Our first objective was to establish a pre-vaccination baseline for VT carriage prevalence among college students as it will allow us to assess the impact of MenACWY vaccine in the Netherlands in the future. The second objective was to investigate the use of saliva samples to monitor meningococcal carriage.

## MATERIALS and METHODS

### Ethics statement

The study protocol was reviewed by the Centre for Clinical Expertise at the RIVM. Since procedures were considered non-invasive, and participants were anonymized, the study was considered outside the ambit of the WMO (Medical Research Human Subjects Act, www.ccmo.nl). Consequently, the committee approved the consent procedure and granted a waiver for further ethical review.

### Study design and sample collection

In the fall of 2018, saliva and oropharyngeal swabs were collected from college students of Hogeschool Utrecht (n=300). After signing informed consent, students self-collected saliva by spitting 1ml into a 15ml tube (Greiner, Kremsmünster, Austria). Next, a study nurse swabbed student’s posterior pharyngeal wall with a nylon swab (FLOQSwabs, COPAN, Brescia, Italy) to collect an oropharyngeal sample. Immediately after collection, saliva (approximately 50 µl) and oropharyngeal swab were used to inoculate Neisseria Selective Medium PLUS agar plates (NS-agar; Oxoid, Badhoevedorp, the Netherlands) and within 20 minutes plates were placed in a 37°C, 5% CO_2_ incubator. Once all samples have been collected, cultured plates were transported at room temperature to the laboratory.

### Meningococcal carriage detection using culture

Upon arrival, NS-agar cultures were incubated for up to two days at 37°C and 5% CO_2_. On both days cultures were screened for presence of meningococcus-like colonies (grey, round and smooth colonies with convex shape). When found, 1-3 colonies were re-plated on Columbia Blood agar (CBA, bioTRADING Benelux B.V., Mijdrecht, the Netherlands) and tested for species identification using Matrix-assisted Laser Desorption/Ionization Time-of-Flight mass spectronomy (MALDI-ToF; Bruker Daltonik GmbH, Bremen, Germany).

Separately for OP and saliva samples, a single isolate with a score ≥2.0 for *Neisseria meningitidis* (database BDAL V8.0.0.0+SR1.0.0.0, Bruker Daltonik) was stored at −70°C in Brain Heart Infusion (BHI, Oxoid) supplemented with 0.5% Yeast Extract (YE; Oxoid) and 10% glycerol. NS-agar cultures displaying any microbial growth were harvested into 2 ml of Todd-Hewitt Broth (Oxoid) supplemented with 0.5% YE and 10% glycerol. These harvests were considered to be culture-enriched for meningococci, and 0.7 ml of it stored at −70°C.

### Detection of meningococcal DNA with qPCR

DNA was extracted from 100 µl of harvest of culture-enriched samples using DNeasy Blood & Tissue kit (Qiagen, Hilden, Germany) as previously described [22]. DNA eluted into 100 μl sample volume was tested in quantitative-PCRs (qPCRs) using primers and probes (Eurogentec, Seraing, Belgium) targeting sequences within *metA*, a gene encoding for a periplasmic protein, and a capsule transporter gene *ctrA* [23, 24]. The qPCRs were conducted using Probes Master 480 (Roche) mastermix, primers and probes concentrations are listed in **Table S1**, with 2 μl of DNAused in 12.5 μl reaction volumes. The qPCR assays were conducted on LightCycler480 (Roche) with programme as described in **Table S2**. A 10-fold serial dilution of DNA from a meningococcal strain (**Table S3**) was used as standard curve. C_T_ tresholds for positivity were determined with Youden index calculated using Receiving operating characteristic (ROC) curve analysis [25].

### Recovery of live meningococcu*s* from culture-enriched samples

To test whether lower sensitivity of conventional diagnostic culture could account for differences between qPCR and culture results, culture-enriched samples first classified as negative by culture yet positive by qPCR were revisited to recover viable meningococci. For this second culture guided by qPCR results, CBA plates were inoculated with 100 μl culture-enriched sample in 10^−2^-10^−4^ dilutions, incubated at 37°C and 5% CO_2_, and screened for meningococcus as decribed above.

### Genogroup-specifc qPCRs

Two microliters of DNA extracted from culture-enriched samples were tested in 12.5μl of reaction volume in qPCRs targeting genogroups A, B, C, W or Y [24]. Primer and probe concentrations are listed in **Table S1**. These qPCRs were conducted on a LightCycler480, using SensiFast probe No-ROX mastermix (Bioline, London, United Kingdom) and with programme described in **Table S2**. Culture-enriched samples were regarded as positive for a genogroup when the C_T_ was lower than the cut-off value set for *metA* and *ctrA*. Control strains are listed in **Table S3**.

### Genotyping of meningococcal strains

DNA extracted from cultured strains was tested in *metA*, *ctrA* and genogroup-specifc qPCRs. Since not all genogroups were covered by qPCRs, a simplified criterium of positivity for *ctrA* was also applied to classify strain as genogroupable.

### Statistical analysis

Data was analyzed using Prism (GraphPad Software; v8.4.1) and R (version 4.0.0). A *p* value of <0.05 was considered significant. ROC curve analysis was performed using “cutpointr” package [25], and Cohen’s Kappa (*κ*) was determined in analysis of methods agreement. Youden index values were determined via bootstrapping (n=1,000) on *metA* qPCR data from saliva and OP to determine the optimal cut-off value for qPCR detection [25].

## RESULTS

All samples were collected in October and November 2018. Of 300 students that consented to participate, one person refused to have the oropharynx swabbed and was excluded from the study. Paired saliva and oropharyngeal samples from the remaining 299 (61% female; median age 20 years, range 16-28 years) students were analyzed (**Figure 1**).

**Figure 1:**
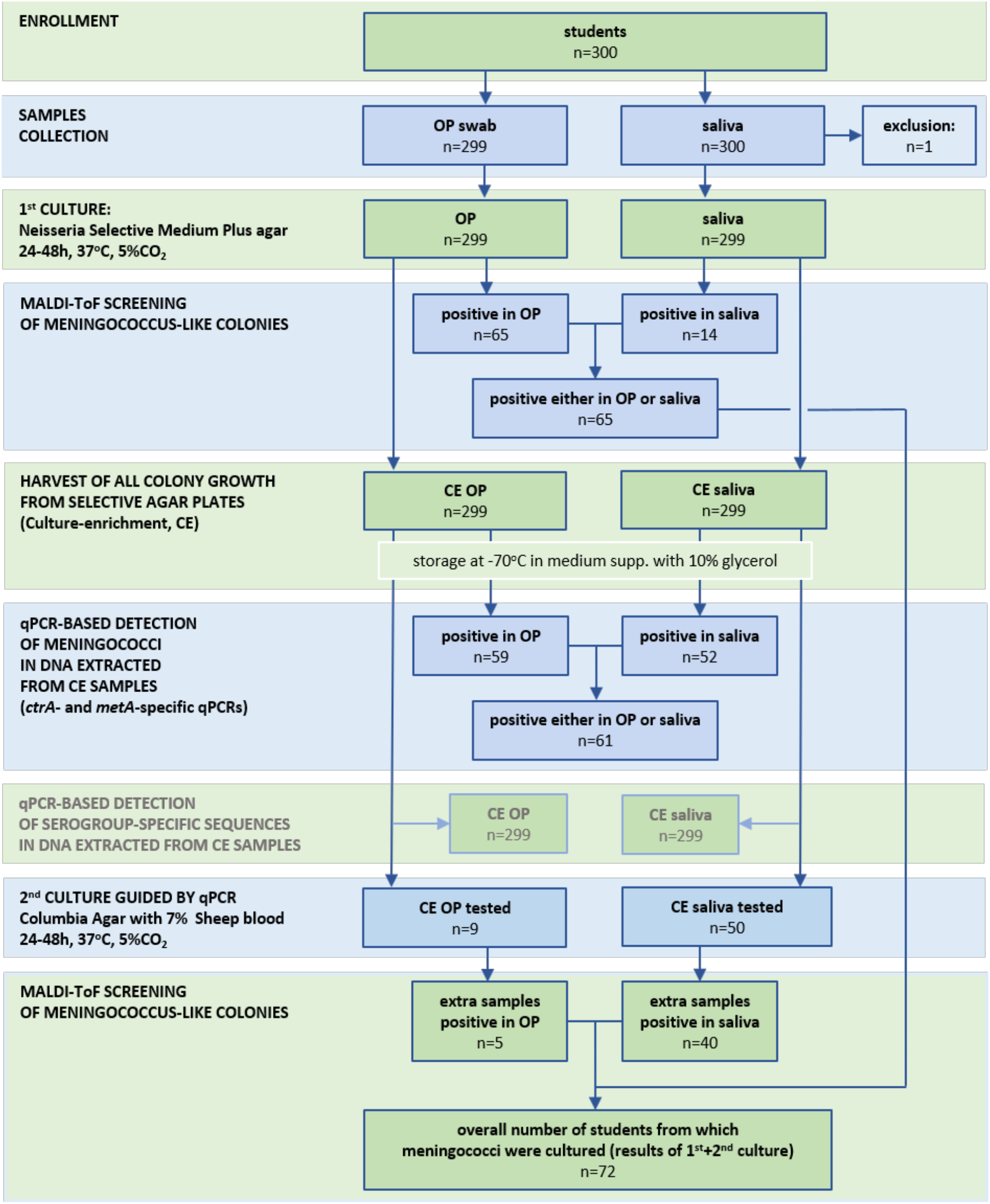
Flowchart depicting the study workflow and results of meningococcal detection using either culture-based or qPCR-based diagnostic methods.

Bacterial strains classified with MALDI-ToF as meningococcus were cultured from 72 students (24% of 299) of which 70 had strains isolated from the oropharynx and 54 from saliva (**Table 1**). Sixty-five (93%) of 70 oropharyngeal samples positive by culture had meningococcus isolated from the first culture and the remaining five strains were recovered when samples positive by qPCR yet initially culture-negative for meningococcus were revisited. For saliva, the same procedure resulted in fourteen samples positive for meningococcal strains in the first culture (26% of 54) and the remaining 40 in cultures guided by qPCR showing that initial diagnostic cultures displayed vastly reduced sensitivity for saliva when compared with oropharyngeal samples (14 vs. 65 strains cultured from 299 students, McNemar’s test, *p*<0.0001). The differences also remained significant after qPCR-guided culture (54 vs. 70, *p*<0.001). Genogroupable meningococci were cultured from 62 students (21% of 299). Here too, the number of culture-positive samples was significantly higher for oropharyngeal swabs compared with saliva (58 vs. 45, *p*<0.01) (**Table 2**).

**Table 1:** The accuracy of *Neisseria meningitidis* detection in oropharyngeal and saliva samples collected from 299 students and tested with culture and using molecular methods applied to DNA extracted from culture-enriched samples. Measures of diagnostic accuracy were calculated by comparing the number of individuals positive per method with the overall number of individuals positive for *N. meningitidis* by any method.

| Method | Oropharyngeal swab |  |  |  |  |  |  | Saliva |  |  |  |  |  |  |
| --- | --- | --- | --- | --- | --- | --- | --- | --- | --- | --- | --- | --- | --- | --- |
|  | Prevalence | PPV | NPV | Sensitivity | Specificity | Concordance | K | Prevalence | PPV | NPV | Sensitivity | Specificity | Concordance | K |
|  | %<br>(95%CI) | % | % | %<br>(95%CI) | % | % |  | %<br>(95%CI) | % | % | %<br>(95%CI) | %<br>(95%CI) | % |  |
| initial culture | 21.7<br>(17.4–26.8) | 96.1 | 97.0 | 99.1<br>(96.5–99.8) | 87.8<br>(78.2–93.5) | 96.3 | 0.90 | 4.7<br>(2.8–7.7) | 78.9 | 100 | 100<br>(–) | 18.9<br>(11.4–29.6) | 79.9 | 0.26 |
| qPCR | 19.7<br>(15.9–25.0) | 93.8 | 100 | 100<br>(–) | 79.7<br>(69.0–87.5) | 95.0 | 0.86 | 17.4<br>(13.5–22.1) | 91.1 | 100 | 100<br>(–) | 70.0<br>(58.8–79.6) | 92.6 | 0.78 |
| initial plus qPCR-guided cultures | 23.4<br>(19.0–28.5) | 98.3 | 100 | 100<br>(–) | 94.6<br>(85.8–98.1) | 98.7 | 0.96 | 18.1<br>(14.1–22.8) | 91.5 | 100 | 100<br>(–) | 71.6<br>(60.2–80.8) | 93.0 | 0.79 |

PPV: positive predictive value, NPV: negative predictive value, 95%CI: 95% confidence interval, $\kappa$ : Cohen's Kappa where $\leq 0$ , 0.01-0.20, 0.21-0.40, 0.41-0.60, 0.61-0.80, $>0.81$ are interpreted as no agreement, none to slight, fair, moderate, strong, and almost perfect agreement, respectively.

**Table 2:**
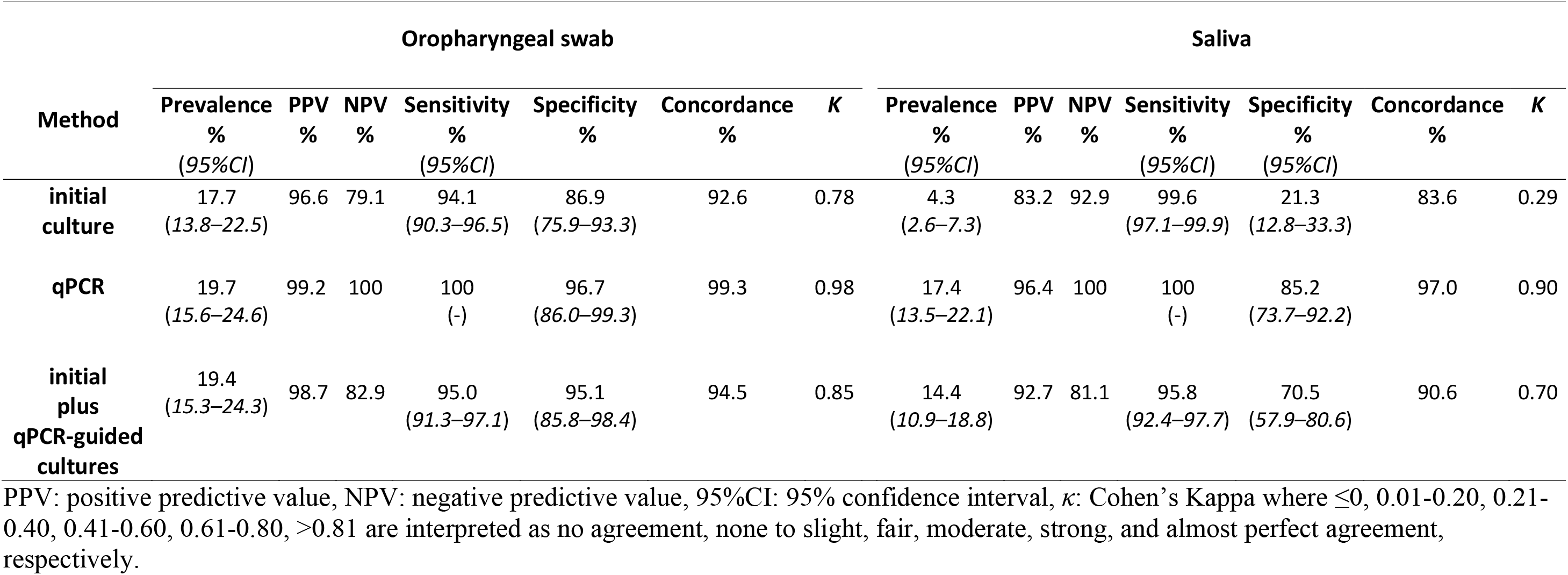
The accuracy of genogroupable *Neisseria meningitidis* detection in oropharyngeal and saliva samples collected from 299 students and tested with culture and using molecular methods applied to DNA extracted from culture-enriched samples. Measures of diagnostic accuracy were calculated by comparing the number of detected individuals positive per method with the overall number of individuals positive for genogroupable *N. meningitidis*.

The study criterium for classification of a sample as positive for *N. meningitidis* by qPCR was detection of both *metA* and *ctrA* in DNA extracted from a culture-enriched sample, and was derived by calculating the optimal C_T_ cut-off values by using the Youden index (**Table S4**). Using qPCR detection we identified 61 students (20% of 299) as a meningococcal carrier. The difference in proportion of carriers detected by qPCR between oropharyngeal samples and saliva samples was not significant (59 or 20% vs. 52 or 17%, McNemar’s test; *p*=0.07), both methods showed high agreement (96%; *κ*=0.88). Samples classified as positive for meningococcus by qPCR displayed significant correlation between *metA* and *ctrA*, supporting high specificity of molecular detection (**Figure 2**). Detection by culture and by qPCR resulted in 71 (24% of 299) and 62 (21%) students identified as a meningococcal carrier in oropharyngeal and saliva samples, respectively. Altogether 74 (25%) students were identified as a meningococcal carrier and 62 (21%) as carrier of genogroupable meningococci by any method used (**Figure 3**).

**Figure 2:**
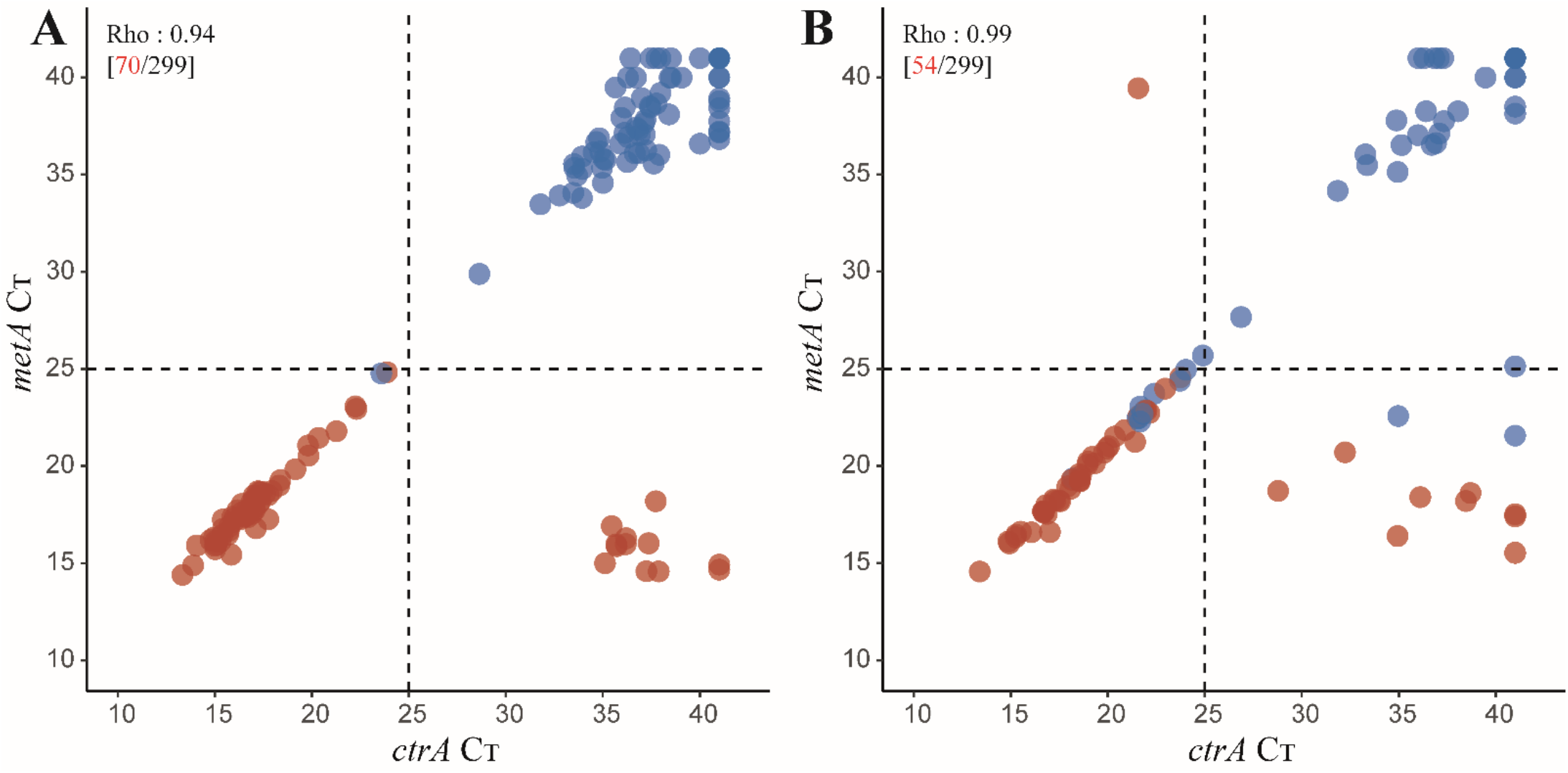
qPCR based detection of *Neisseria meningitidis* versus isolation of live meningococci from oropharyngeal and saliva samples. A scatter plot of the *metA* and *ctrA* qPCR cycle threshold (C_*T*_) values from (**A**) oropharyngeal and (**B**) saliva samples. Each symbol represents an individual sample. Samples with a C_*T*_ for both *metA* and *ctrA* below 25 C_T_ are considered as positive for meningococcal carriage when tested with molecular methods. In both oropharyngeal and saliva samples, we noted a significant correlation between *metA* and *ctrA* for meningococcus positive samples (Spearman’s test *p*<0.0001). Red dots represent samples from which meningococcal strain has been cultured. Blue dots represent samples classified as positive for meningococcus when tested with molecular method but negative by culture. Numbers in brackets depict the number (in black) of all samples and (in red) number of samples from which *N. menigitidis* has been cultured.

**Figure 3.**
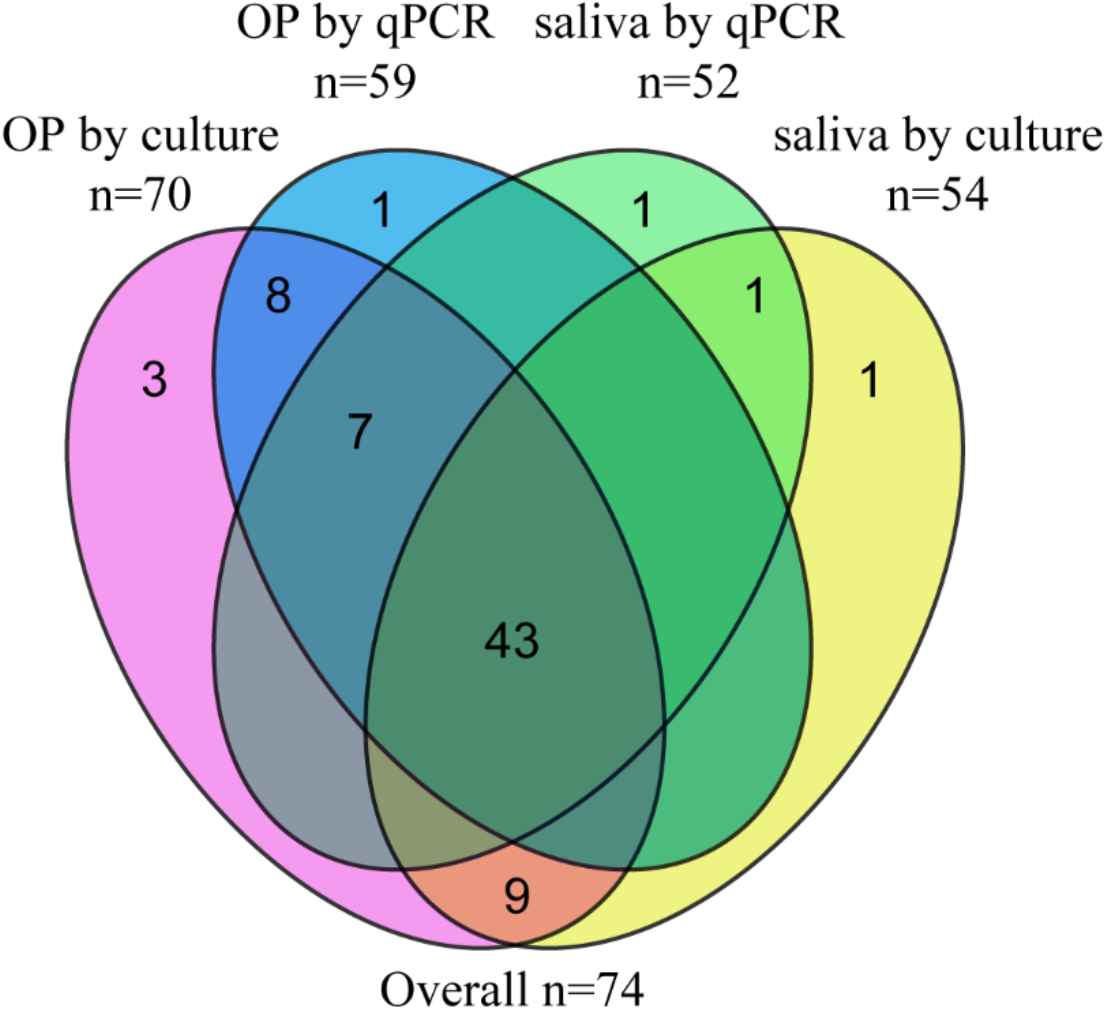
Venn diagram displaying the number of oropharyngeal and saliva samples positive for meningococci based on recovery of live *N. meningitidis* strain from a culture (samples positive by culture, includes qPCR-guided culturing) or when tested with qPCR.

When comparing methods and specimen types used for detection of meningococcal carriage overall, all evaluated procedures displayed comparable specificity of detection (**Table 1**) and primarily varied in performance for sensitivity and for positive predictive value (PPV).

The criterium based on both *ctrA* and *metA* was expected to impact negatively the sensitivity of meningococcal carriage detection by qPCR when compared with culture due to the presence of non-genogroupable meningococci that were likely to be *ctrA*-negative. Therefore, we compared methods and specimen types on samples containing genogroupable meningococci (**Table 2**), which were supposed to be positive for both *ctrA* and *metA*. The PPV and sensitivity of the evaluated methods were highest for detection by qPCR, whereas using saliva samples resulted in decreased negative predictive values (NPV) when compared with oropharyngeal samples. Detection of genogroupable meningococci using qPCR and saliva displayed increased PPV and comparable sensitivity when compared with detection of meningococcus in initial oropharyngeal cultures.

Next, we determined with genogroup-specific qPCRs the prevalence of genogroup A, B, C, W and Y carriage. The specificity of these qPCR assays was tested using culture-enriched samples negative for meningococcal carriage by culture and qPCR and none of the samples negative for *ctrA* generated a signal below 25 C_T_ for a genogroup-specific gene (**Figure S1**). Altogether, 51 (83.6%) of 61 students identified as carriers of meningococci with qPCR were positive for any of the genogroups targeted in group-specific qPCRs (**Figure 4**). Genogroup-specific C_T_s of almost all samples positive for any of the tested genogroups corresponded strongly to the C_T_ for *ctrA*. The exception was a single oropharyngeal sample for which results were indicative of potential co-carriage of a genogroup Y strain with another *ctrA*-positive meningococcal strain of unidentified group (**Figure 4E**). The prevalence of genogroup B (8.7% of 299) and Y (6.4% of 299) was highest while genogroups C (0.7% of 299) and W (1.3% of 299) were less prevalent. None of the samples were positive for genogroup A. MenACWY VT genogroups accounted for 8.4% (95%CI 5.7-12.1) carriage prevalence or 41.0% of meningococcal identified by qPCR. Results between specimen types were concordant for genogroups (**Table 3**).

**Figure 4.**
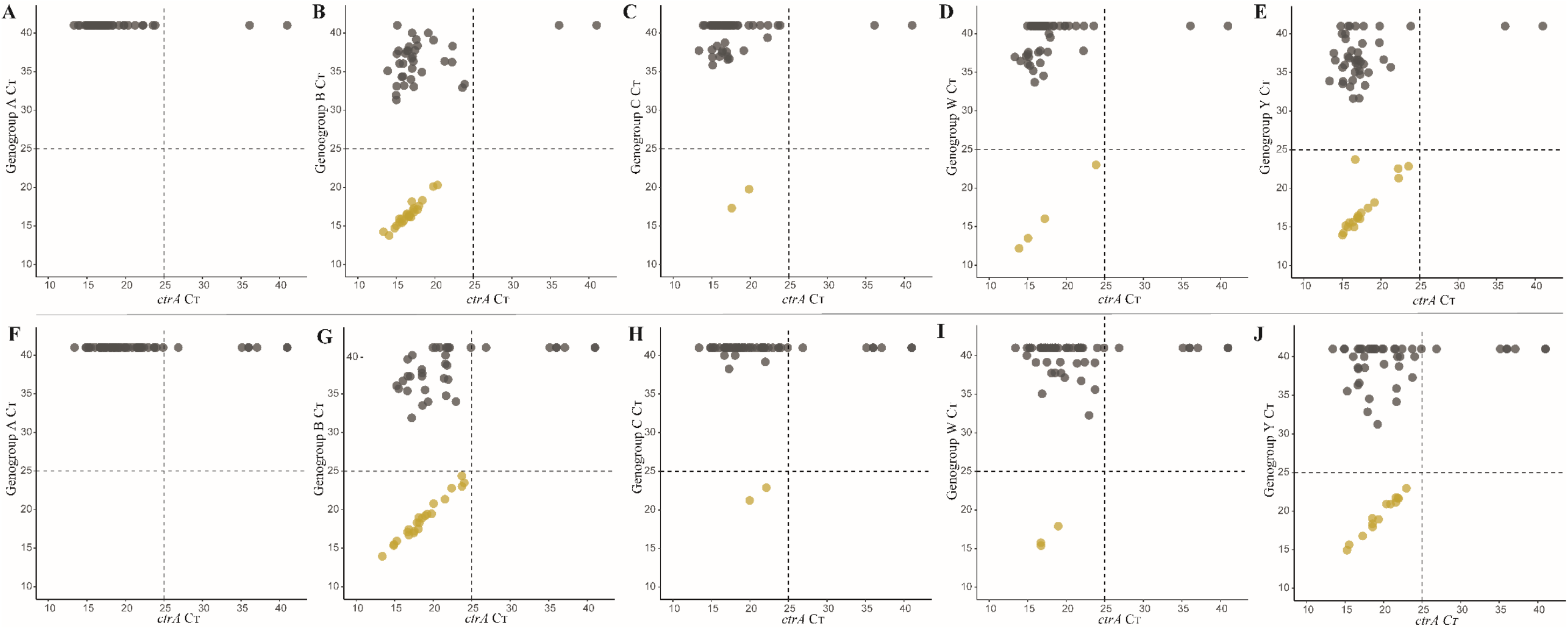
A scatter plot of the *ctrA* and genogroup-specific qPCR cycle threshold (C_T_) values. Results are displayed for oropharyngeal (**A** – **E**) and saliva (**F** – **J**) samples. Each dot represent an individual sample. Samples with a C_T_ for both *ctrA* and a particular genogroup below 25 C_T_ are considered as positive for that particular genogroup. Yellow dots represent samples classified as positive for a genogroup by qPCR and grey dots as negative for the depicted genogroup. Dashed lines depict the C_T_ criterium for meningococcal carriage.

**Table 3.** Prevalence of meningococcal MenACWY vaccine-type serogroups among OP and saliva samples collected from students (n=299) and tested by qPCR.

| Parameter | OP<br>n (%)<br>(95% CI) | Saliva<br>n (%)<br>(95% CI) | OP and saliva<br>n (%)<br>(95% CI) | Either OP or saliva<br>n (%)<br>(95% CI) | Concordance | P value* |
| --- | --- | --- | --- | --- | --- | --- |
| menA | 0 | 0 | 0 | 0 | - | - |
| menB | 25 (8.3)<br>(5.2-11.3) | 23 (7.7)<br>(5.2-11.3) | 22 (7.3)<br>(4.9-11.0) | 26 (8.7)<br>(6.0-12.4) | 98.7% | 0.6171 |
| menC | 2 (0.7)<br>(0.1-2.4) | 2 (0.7)<br>(0.1-2.4) | 2 (0.7)<br>(0.1-2.4) | 2 (0.7)<br>(0.1-2.4) | 100% | - |
| menW | 4 (1.3)<br>(0.5-3.4) | 3 (1.0)<br>(0.3-2.9) | 3 (1.0)<br>(0.3-3.0) | 4 (1.3)<br>(0.5-3.4) | 99.7% | 1.0000 |
| menY | 18 (6.0)<br>(3.8-9.3) | 14 (4.7)<br>(2.8-7.7) | 13 (4.3)<br>(2.6-7.3) | 19 (6.4)<br>(4.1-9.7) | 98.0% | 0.2207 |

\*: *p*-values are calculated from McNemar tests comparing students positive for serogroup in oropharyngeal samples and saliva samples. The percentage of concordance displays the proportion of samples (n=63) with identical result in serogroup-specific qPCR assay for a particular serogroup.

## DISCUSSION

In this cross-sectional study, we evaluated the application of saliva for meningococcal carriage detection using both culture and qPCR-based methods. Our goal was to optimize meningococcal detection for future carriage studies assessing the impact of the meningococcal vaccines on carriage. Although meningococcal detection with culture resulted in fewer meningococcal carriers identified in saliva when compared with oropharyngeal samples, qPCR detection of meningococcus did not result in significant differences between these two sample types.

Based on culture, we observed an overall carriage prevalence of 24.1%. The difference in positivity for meningococcus between OP and saliva samples was likely caused by a greater difficulty to isolate meningococci from saliva cultures. In the saliva, a higher abundance of commensal species capable of growth on the culture media was observed as described previously [18]. Using qPCR detection we observed an overall carriage prevalence of 20.4%, and no significant differences were observed between OP and saliva samples in positivity for meningococcus. Importantly, when detecting carriage of genogroupable meningococci, the method of testing culture-enriched saliva with a qPCR performed at least equally well compare with conventional diagnostic culture of oropharyngeal swab. Although numerous studies have implicated oral fluids in meningococcal transmission [8–16], very few actually tested saliva as specimen for assessing meningococcal carriage [18, 19]. In this context, our findings are in line with a meningococcal carriage study conducted recently by Rodrigues *et al*. [20].

Among 299 students, we observed an overall meningococcal carriage rate of 24.7%, a prevalence that is in line to what has been reported previously with pharyngeal swabs for this age group [2, 15]. The prevalence of meningococcal carriage among young adults is considered to be higher than other age groups due to increased social interactions which facilitate meningococcal transmission [6]. In addition, age-related alterations in the microbiota of the URT may prime individuals for meningococcal colonization [26].

VT serogroups targeted in the MenACWY vaccine accounted for 41.0% of meningococci detected in carriage, corresponding to a prevalence of 8.4%. Of these VT genogroups, genogroup Y was most frequently detected. While an outbreak of serogroup W was ongoing in the Netherlands during the fall of 2018, the prevalence of genogroup W in carriage was low (1.3%). The prevalence of genogroup C was also low, possibly reflecting reduced circulation since implementation of menC vaccine in the Netherlands [27]. Serogroup A was not detected in our study, its circulation appears to be limited in the Netherlands [15, 28]. The most prevalent genogroup among carriers was B. Genogroups B and Y have both been described to be most commonly detected genogroups among young adults [29].

Our study has a number of limitations. Firstly, MALDI-ToF may have identified more samples of students positive for meningococcus than qPCR detection with *metA* and *ctrA* carriage criterium as MALDI-ToF also takes non-genogroupable, unencapsulated meningococci into account, and is susceptible to misidentification [30]. To avoid misidentification byby MALDI-ToF, we have only included bacterial strains for which identification displayed high confidence (≥2.0). Another limitation is false-positivity of qPCR tests. To minimize this issue, we have conducted ROC curve analysis and used the Youden index to determine a cut-off value for qPCR detection. Considering that the majority of qPCR positive samples facilitated successful recovery of viable meningococci, we conclude that false-positive results have had no significant impact on our conclusions.

One of the strengths of our study was the paired comparison of saliva and oropharyngeal samples in detection of meningococcal carriage. Furthermore, we have used selective media and inoculated plates immediately after samples collection. Fast processing of samples may be crucial for the sensitivity of meningococcal detection. Moreover, the combined use of two meningococcal qPCR targets for specific meningococcal detection in polymicrobial samples has allowed us to detect meningococcus with high specificity.

In conclusion, our findings show that the detected prevalence of meningococcal carriage between oropharyngeal and saliva samples was nondifferent with qPCR detection, the results for saliva were highly concordant with oropharyngeal swabs, and that the majority of samples positive with qPCR were shown to contain viable meningococci. Since the collection of saliva is easy, well tolerated and can be performed without professional assistance, we propose that saliva combined with qPCR-based surveillance can be considered for future meningococcal carriage studies.

## Authors’ contribution

EAMS, TB and KT had an idea and initiated the study. AJWM, NY and TB secured financial support for the project. TB and KT led the project and supervised the project activities. WRM, AJWM, NYR, TB and KT wrote the protocol. WRM, GP, LJdJ, IG, SW, and JvV validated the methods. WRM, GP, LJdJ, IG, SW, and EvL conducted the research and collected the data. WRM, RM, MMI, AJW, NYR, TB and KT managed the study. WRM, RM, GP, and KT curated the data. WMR, RM and KT performed formal analysis of study data. WRM and KT visualized presentation of the results and drafted the manuscript. All authors amended, critically reviewed and commented on the final manuscript.

## Acknowledgements

We gratefully acknowledge the students of Hogeschool Utrecht for their participation in the study. We wish to thank Carien Voogt for assistance in setting-up molecular assays. This study was presented in part at ECCMID 2019 in Amsterdam, the Netherlands.

---

**Supplementary Table S1:**
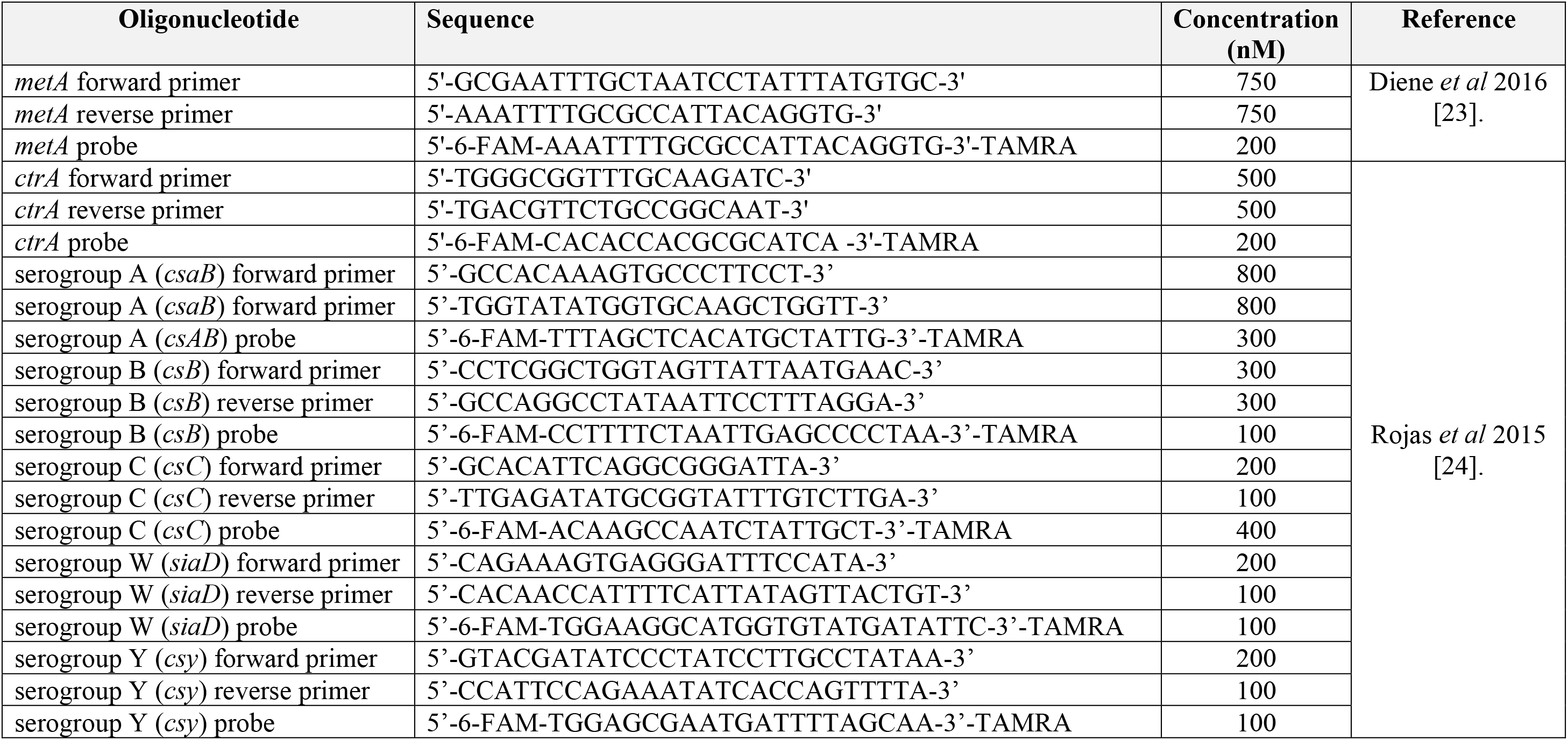
Primer and probe concentrations used in the study.

**Supplementary Table S2:**
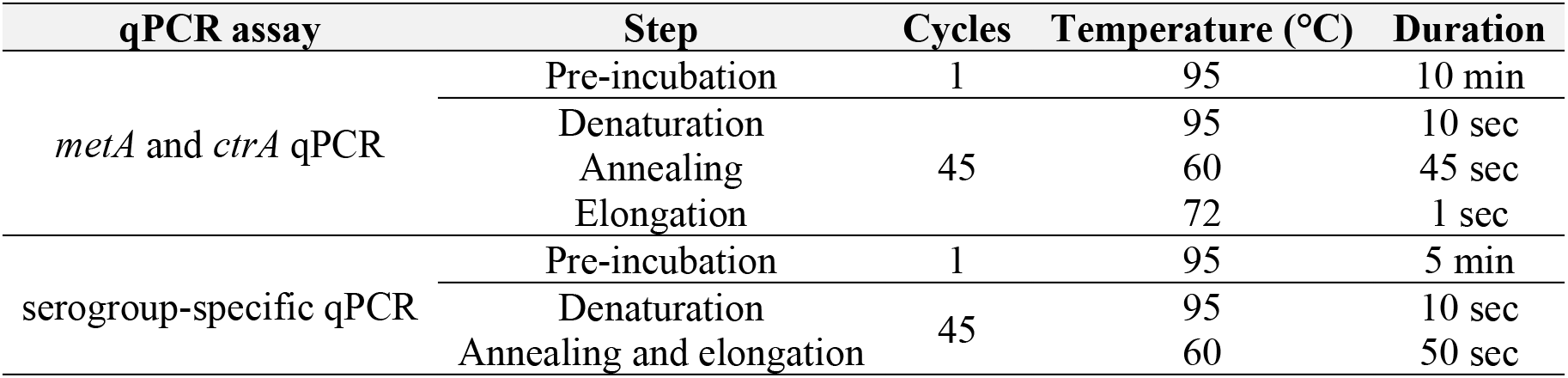
qPCR programmes used in this study.

| qPCR assay | Step | Cycles | Temperature (°C) | Duration |
| --- | --- | --- | --- | --- |
| *metA* and *ctrA* qPCR | Pre-incubation | 1 | 95 | 10 min |
|  | Denaturation | 45 | 95 | 10 sec |
|  | Annealing |  | 60 | 45 sec |
|  | Elongation |  | 72 | 1 sec |
| serogroup-specific qPCR | Pre-incubation | 1 | 95 | 5 min |
|  | Denaturation | 45 | 95 | 10 sec |
|  | Annealing and elongation |  | 60 | 50 sec |

**Supplementary Table S3:** *Neisseria meningitidis* strains used in this study.

| Strain | Description | Source |
| --- | --- | --- |
| serogroup A strain 3125 | used to optimize the serogroup-specific qPCR assay | Meningococcal Reference Unit Manchester, UK |
| serogroup B strain BD00-00032 | used to optimize the serogroup-specific qPCR assay | this study |
| serogroup C strain BD00-00268 | used to optimize the serogroup-specific qPCR assay | this study |
| serogroup W strain BD98-00112 | used to optimize the serogroup-specific qPCR assay, *ctrA* qPCR assay and *metA* qPCR assay. | this study |
| serogroup Y strain BD03-00373 | used to optimize the serogroup-specific qPCR assay | this study |

**Supplementary Table S4:** Optimal qPCR C_T_ threshold and corresponding parameters for meningococcal carriage detection on samples stratified by positive or negative for culture detection. For qPCR detection of *Neisseria meningitidis*, we regarded a culture-enriched sample as positive by qPCR when detection of both the *metA* and *ctrA* genes was observed. For both types of samples and in both qPCRs we observed a bimodal distribution of C_T_s, with the highest C_T_ of any culture-positive sample separated by at least 10 C_T_s from the lowest in a cluster of all culture-negative samples. Based on this distribution, we performed ROC curve analysis to calculate the maximal Youden indices. For oropharyngeal samples, the difference between thresholds for positivity calculated for *metA* and *ctrA* was within 0.1 C_T_. For saliva samples the diference was over 13 C_T_s due to higher than among oropharyngeal samples proportion of non-genogroupable to groupable strains cultured. To avoid a bias in meningococcal detection between oropharyngeal and saliva samples, we applied thresholds (<25 C_T_) calculated for oropharyngeal samples also to saliva. A criterium based on both *ctrA* and *metA* was expected to impact negatively the sensitivity of meningococcal carriage detection by qPCR when compared with culture due to presence of non-genogroupable meningococci that were likely to be *ctrA*-negative.

| Parameter | Optimal threshold | Youden index (J) | Sensitivity | Specificity |
| --- | --- | --- | --- | --- |
| **OP *metA*** | 24.83 | 0.9956 | 1 | 0.9956 |
| **OP *ctrA*** | 24.87 | 0.8242 | 0.8286 | 0.9956 |
| **Saliva *metA*** | 24.95 | 0.9407 | 0.9815 | 0.9592 |
| **Saliva *ctrA*** | 38.70 | 0.8302 | 0.9444 | 0.8857 |

**Figure S1.**
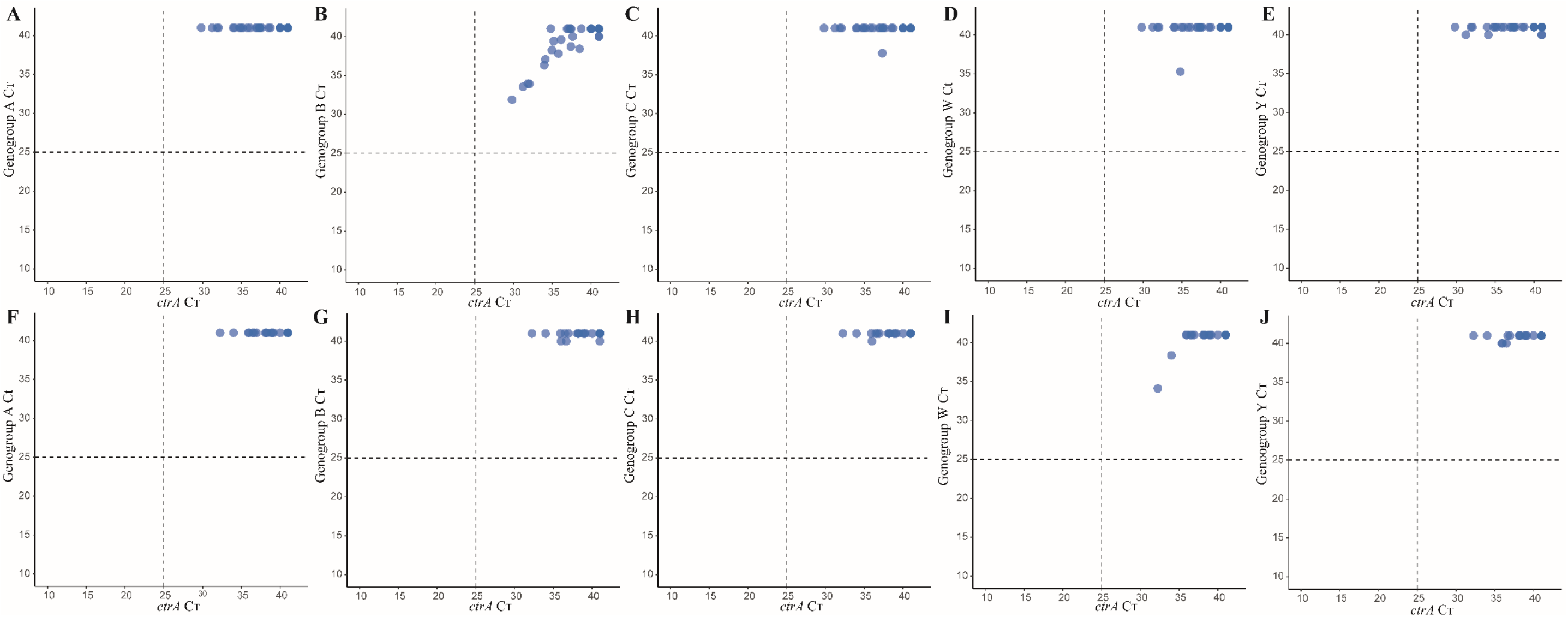
Scatterplots of genogroup-specific qPCR assays for CE samples negative for meningococcus by qPCR. Scatterplots displays genogroups-specific qPCR results for culture-enriched oropharyngeal (**A** – **E**) and saliva (**F** – **J**) samples negative for meningococcal carriage by qPCR and culture (n=42 for each). None of the tested samples generated a signal (C_T_) below 25 C_T_ for any of the tested genogroups, namely serogroup A, B, C, W and Y. Dashed lines depict the C_T_ criterium for meningococcal carriage.

